# QKI 6 ameliorates CIRI through promoting synthesis of triglyceride in neuron and inhibiting neuronal apoptosis associated with SIRT1-PPARγ-PGC-1α axis

**DOI:** 10.1101/2020.12.23.424095

**Authors:** Rui Liu, Hongzeng Li, Jingyuan Deng, Qunqiang Wu, Chunhua Liao, Qun Xiao, Qi Chang

## Abstract

The stroke induced by ischemia of brain remains high incidence and death rate. The study wanted to confirm the effects of QKI 6 on the protection role in neurons of rat model of cerebral ischemia/reperfusion injury (CIRI). The rat model with CIRI induced by MCAO (middle cerebral artery occlusion) was well established and rat neurons were isolated to characterize the effects of QKI 6 mediated by SIRT1 on synthesis of triglyceride in neuron and neuronal apoptosis via activation of SIRT1-PPARγ-PGC-1α signaling pathway. The expression levels of SIRT1 or QKI 6, and acetylation level of QKI 6 was decreased in neurons of rat model with CIRI. QKI 6 deacetylated and mediated by SIRT1 that contributed to suppressing the progression of neuronal apoptosis in rat through promoting synthesis of triglyceride *in vivo* and *in vitro* via SIRT1-PPARγ-PGC-1α signaling pathway, then inhibiting CIRI. In conclusion, our results demonstrated SIRT1 deacetylates QKI 6, the RNA-binding protein, that affects significantly the synthesis of triglyceride in neurons of CIRI rat model. Moreover, it activated transcription factor PGC-1α through post-transcriptional regulation of the expression of PPARγ, and further enhanced synthesis of triglyceride, thereby restrained the progression of neural apoptosis and CIRI.

## Introduction

The stroke induced by ischemia of brain remains high incidence and death rate, and is the considerable reason of eternal death and disability. Therefore, ischmic stroke can cause the obvious burden of community economy and health [1]. CIRI (Cerebral ischemia/reperfusion injury) is induced by hypoxia and ischemia of brain, and the usual course related with the pathological status of ischemic stroke. Moreover, in the area of ischemia, CIRI further accentuated by blood flow reperfusion [2]. The specturm of complex pathogenesis in CIRI includes inflammation, calcium dysregulation, excitotoxicity, apoptosis, and necrosis. These are a great deal of neurological injury, which can make finally for the irreversible tissues damage in brain [3]. CIRI is also the key health problem and challenge, though ischemic stroke is treated with regimens based on the wonderful advances [4, 5]. Therefore, the refined mechanism of CIRI and the effective therapy strategy of ischemic stroke needed to be identified with more efforts. Many previous investigations want to explore the effective therapies or medicines for prevent and alleviate ischemic damage [6, 7]. Intriguingly, based on the different of anti-inflammation, anti-apoptosis, and anti-excitotoxicity mechanisms, CIRI can be relieved in opposition to cerebral ischemia with neuroprotective effects induced by several drugs [8].

Through regulating the protein modification, SIRT (Sirtuins) provide the spectrum of bio-functions by deacetylating cellular proteins at post-translational level [9]. From bacteria to humans, the sirtuins are widely conserved deacetylased dependent on NAD (nicotinamide adenine dinucleotide), and regulate a series of functions in central neural system (CNS). The obvious evidences demonstrate that SIRT1 maybe a potential strategy for the therapy of neuro-degenerative diseases. The various of neurological disorders and traumatic injury can induce neuronal degeneration usually accompanied with inflammation. Moreover, SIRT1 activation can significantly suppress inflammatory responses, and is the sole therapy for controlling both inflammatory cells and neurons [9].

QKI have different bio-functions in relation to the stability [10–12] or translation [13, 14] of mRNA, the processing of miRNA [15, 16], even selective splicing [14, 17–21]. We have recently found that SIRT1 increased QKI 6 acetylation level by suppressing the activity of SIRT1. In contrary, the increasing activity of SIRT1 caused deacetylation of QKI 6. Either in neurons knockdown of SIRT1 or neural tissues of SIRT1-silenced rat, the increased level of QKI 6 acetylation presented. Our findings implicated the selective splicing in neurons maybe the crucial factor in regulation of lipid metabolism.

Lipid metabolism is associated with a great deal of cellular processes, such as proliferation, differentiation, growth, survival, apoptosis, motility, inflammation, membrane homeostasis, drug resistance, and chemotherapy response. Moreover, bioactive lipid molecules can enhance cell intrinsic apoptotic pathway through activating caspases enzymes and regulating the permeability of mitochondrial membrane. To understand and define the lipid metabolism of neurons is very important to explore the novel strategy for anti-stroke therapy [22].

Therefore, the role of QKI 6 mediated by SIRT1 in this work will be elucidated on the pathological process in CIRI model. Ischemic stroke caused by the the MCAO (Middle cerebral artery occlusion) model subsequently combined with reperfusion [23]. The CIRI rat model induced by MCAO was well established and neurons of rats were segregated for identifying the functional characterization of QKI 6 associated with SIRT1 and the activation of PPARγ/PGC-1α signaling pathway on the area of neuronal apoptosis in CIRI rat model.

## Materials and methods

### Primary culture for cerebral cortical neurons of rat

Our experiments were performed according to Guidelines for use of Laboratory Animals in Fourth Military Medical University (Xi’an, China). Based on previous methods come from Oka et al [24], cerebral cortical neurons of fetal Sprague-Dawley (SD) rats were isolated at 17-18 day-old embryonic rats.

### Oxygen/glucose deprivation (OGD)

In our work, ischemia was induced with OGD [25–27]. The cytosine arabinoside was used to halt the non-neuronal cell division after 4-5 DIV (days *in vitro*). Then, at the DIV of 14-15, mature cultures were performed for the experiments. To initiate ischemia, Hypoxic Workstation incubator was used to change the original one.

### QKI 6 treatment

The protective role induced by QKI 6 were evaluated. before exposing to OGD, primary cortical neurons treated with OGD were pretreated with 50, 100, or 200 ng/mL QKI 6 protein, respectively. Meanwhile, the effect of SIRT1 in the protective role caused by QKI 6 was assessed with SRT1720 (a SIRT1 agonist, 50 ng/mL) and Niacinamide (Nic, a selective inhibitor of SIRT1, 10 μM), Vehicle was used with DMSO. Then cell viability was determined for these cultured neurons with regular medium under normoxia after OGD treatment for another 24 h.

### Examination injury of neurons

The neuronal injury was quantity evaluated based on LDH activity assay [26]. After removing the debris of neurons firstly, the medium of cultured neurons was centrifuged, and then mixed with reagent of the LDH kit for determine with spectrophotometer at 340 nm.

### Determine death of neurons

The death of neurons was analyzed using TUNEL staining and MTT assay. In MTT assays, neurons were dealt with MTT for 2 h at 37°C, and then released formazan of these neurons was quantified at 560 nm using plate reader. Subsequently, the TUNEL stained apoptotic neurons were observed with fluorescence microscope after paraformaldehyde and permeabilized in Triton X-100.

### Analyze the activation of caspase 3

The activations of caspase 3 in neurons were detected with the Activity Assay Kit. Based on manual for the kit, the activation of caspase-3 was examined using fluorescence spectrophotometer at λ_ex_400 nm and λ_ex_505 nm.

### Determination of total TG in primary cerebral cortical neurons of rat

The primary cerebral cortical neurons of rats were cultured in plate after treating with QKI 6 protein. The absorbance at 570 nm of the cell lysate were counted based on standard curve. Then TG concentrations were normalized with protein concentrations. TGs and cholesterol in the primary cerebral cortical neurons were determined with the infinity TC and TGs kit (TR-22421, TR13421; Thermo Fisher Scientific, USA), respectively.

### Analyse QKI 6 acetylation modification induced by SIRT1

KA-predictor (http://sourceforge.net/p/ka-predictor), which was a online tool. The modified sites of acetylation in QKI6 were predicted with the KA-predictor.

### Adenovirus Vector

Ad-SIRT1, the recombinant adenovirus vector was produced using AdEasy (Ad) Vector System (Stratagene, La Jolla, CA, USA). The infection controls were Ad-GFP.

Adenovirus-bacterial recombination system (AdEasy) was used, include pGEM-3ZF (+) and pAdEasy-1 pShuttle-SYN, which was all expression vectors labeled using GFP (green fluorescent protein). The total RNA extracted from cells of rat brain neurons was used for gene amplification of QKI 6, and then it was cloned into vectors based on rat QKI 6 (AF142419.1) sequence in GeneBank. Moreover, the sequence of primer at upward was 5’-atgcttagtctcagcagcctccgcc-3’ and the sequence of primer at downward was 5’-gaccgagccgccaccggcaactaa-3’, QKI 6 was cloned from primary cultured cortical neurons. The evaluation for concentration of virus and viral vectors titer were performed, and then viral stocks were detected on the replication competent viruses with potential contamination. The concentrations of virus were measured at A260 [21]. The titers of viral vectors Ad-QKI 6 and Ad-GFP was 2.5× 10^11^ pfu /ml and 2.0×10^11^ pfu/ml, respectively.

### Cell transfection and siRNA of SIRT1

Followed without or with SRT1720 (50 ng/ml), the neurons treated with OGD were followed with treatment of either scrambled or SIRT1 siRNA. The siRNA of scrambled sequence and SIRT1 siRNA sequence were indicated below: scrambled siRNA: 5’-ACUUUGCUGUAACCCUGUAdTdT-3’ (sense), 5’-UACAGGGUUACAGCAAAGUdTdT-3’ (antisense); SIRT1 siRNA: 5’-CCUACGCCACCAAUUUCGU-3’ (sense), 5’-ACGAAAUUGGUGGCGUAGG-3’ (antisense).

### Immunoprecipitation (IP) with acetylated antibody

The total protein of primary cultured cortical neurons induced by OGD were extracted, then combined with Acetyl Lysine Antibody. After being mixed and incubated overnight at 4°C. The washed sediments were eluted and mixed with the Elution buffer. After centrifuging, the IP products were assessed using western blot assay with the primary antibody of QKI 6 to determine the content of QKI 6 in the products of IP.

### Animals and CIRI model caused by MCAO

Our experiment was approved by Institutional Animal Care and Use Committee of Fourth Military Medical University. The SD rats (250-300 g, adult, female) obtained from Beijing Experimental Animal Center (Beijing, China) were used in our experiment. In standard cages, these rats freed with water and food were housed at 22 ±2°C and cycle of light/dark (12 h/12 h). Ischemic stroke was induced by MCAO with subsequent reperfusion [23]. Rat were anesthetized with atropine (50 mg/kg). At first, the right carotid artery was inserted a nylon filament (diameter, 0.235 mm). After 2 h of advanced occlusion, it was taken out, then blood flow was resupplied with enough reperfusion. The sham rats did not undergo the occlusions but surgery of MCA. After MACO and reperfusion, rat was sacrificed by decollation, brains tissues and infarct volume were collected.

### Adenovirus Infection

Four groups were divided averagely from the 24 rats: Control, Model (CIRI), rat with CIRI and treated with Ad-QKI 6 (QKI 6 group) or vehicle (empty adenovirus vector) (Vehicle group). The group of Ad, Ad-QKI 6 and Vehicle were injected with saline+adenovirus (empty vector), recombinant adenovirus with QKI 6 or saline through the right lateral ventricle for three days before ischemia, respectively. The site of injection based on the brain stereotactic atlas for rat [27]. After continuous administration for 8 weeks, rat were sacrificed. Additionally, brain tissues were subjected to pathological hematoxylin-eosin (HE) and immunohistochemical (IH) stain.

### Measurement of infarct volume and neurological deficit

After reperfusion for 24h, the functions of neurology were assessed with Zea-Longa score [25–27]. The volume of infarction was detected using TTC staining. The sacrifice was conducted after reperfusion, brains were collected and cut into sections (1.0 mm). The sections were fixed with paraformaldehyde after TTC incubation, then imaged for count of infarct volume using Image Pro Plus (version 16.0).

### Western blot

The concentrations of total protein extracted from primary cultured cortical neurons or tissues of brain were quantified using BCA (Bicinchoninic acid assay). Then protein was electrophoresised with SDS-PAGE gel (12%), further transferred onto polyvinylidene difluoride membranes. TBS containing non-fat milk was used to block the membranes, then which was incubated with SIRT1 antibodie (1:800; Cell Signaling Technology, Inc., Danvers, MA, USA), QKI 6 (1:2000; Abcam, Cambridge, UK), PGC-1 α (1:1000; Abcam, Cambridge, UK), PPAR γ(1:1500; Abcam, Cambridge, UK), or anti-cleaved caspase-3 (D175; 1:1200; Cell Signaling Technology, Beverly, MA), β-Actin Mouse mAb (8H10D10; CST #3700; 1:1000; Cell Signaling Technology, Beverly, MA), respectively. Then, those were further incubated using the HRP (horseradish peroxidase)-conjugated second antibodies (1:1000 dilution; Cell Signaling Technology). In the end, the image of blots was obtained with electrochemiluminescence detection system.

### GFP fluorescence detection

The pentobarbital was used terminally to anesthetize the MCAO model after recovery period, brains were made into 2-mm thick coronal sections and sliced from the frontal pole after transcardial perfusion with saline. Then, the sections were fixed with 4% formalin and stained with TTC. The area of infarction in each section was analyzed using the software of SigmaScan Pro 5. The signals of GFP were directly obtained using fluorescence microscope of Olympus BX51. The GFP fluorescent cells in each five view fields selected randomly were counted from one hundred random cells. Since the transfection efficiency was calculated and analyzed using GraphPad Prism 5.0 and Image-ProPlus 6.0.

### Analyzed for NDS (neurological deficit score)

The system of score with modification was used to evaluate neurological functional deficit in the MCAO model after recovery period. NDS of 0 to 10% was considered normal, and 100% was considered maximum deficit.

### Valuation for volume of infarction area

The tissues of brain in MCAO model were firstly sectioned, and then the volume of infarction area was valued with TTC method. The captured image obtained with fluorescence microscope Olympus BX51 (magnification, ×200). In each slice, the infarction area was assessed using image analysis software SigmaScan™ Pro V5.0 and Photoshop 7.0.

### TUNEL staining

At 24 h following MCAO, after fixing with 4% formalin, the brain tissue sections were stained with TUNEL. Five fields of view in the brain tissues sections from contralateral hemisphere were selected randomly. Subsequently, the TUNEL-stained cells were counted, while the digital photograph was observed from slide of brain tissue. In the end, the obtained NDS and apoptotic index were calculated automatically with GraphPad Prism 5.0.

### Statistical analysis

The continuous variables with normally distribute were indicated with mean ± SD (standard deviation). The analysis of statistics was conducted with GraphPad Prism (version 5.0). Student’s *t*-test for comparison of two groups and one-way analysis of variance (ANOVA) with Kruskal-Wallis post hoc tests for multiple group comparisons were performed. *P*<0.05 was statistically difference with significance.

## Results

### QKI 6 played a protective role in neuronal death come from OGD

At first, the protective effects of QKI 6 on OGD-induced death of neurons were determined. The results of MTT experiment indicated that the viability of neurons after treatment with OGD obviously decreased, the survived cortical neurons was merely approximately 55%. However, when neurons were pretreated for 48 h with QKI 6 (50, 100, and 200ng/mL, respectively) before 24h exposure to OGD. The viability assay demonstrated that neurons treated with QKI 6 was remarkably better than control neurons and dose dependence (*P*<0.01) (**Fig. 1A**). Further trial results form leakage rate of LDH assay confirmed that it increased obviously after OGD (**Fig. 1B**). These consequences figured out LDH release could be caused by OGD, which could be significantly inhibited by QKI 6 and dose dependence, too (*P*<0.001).

**Figure 1.**
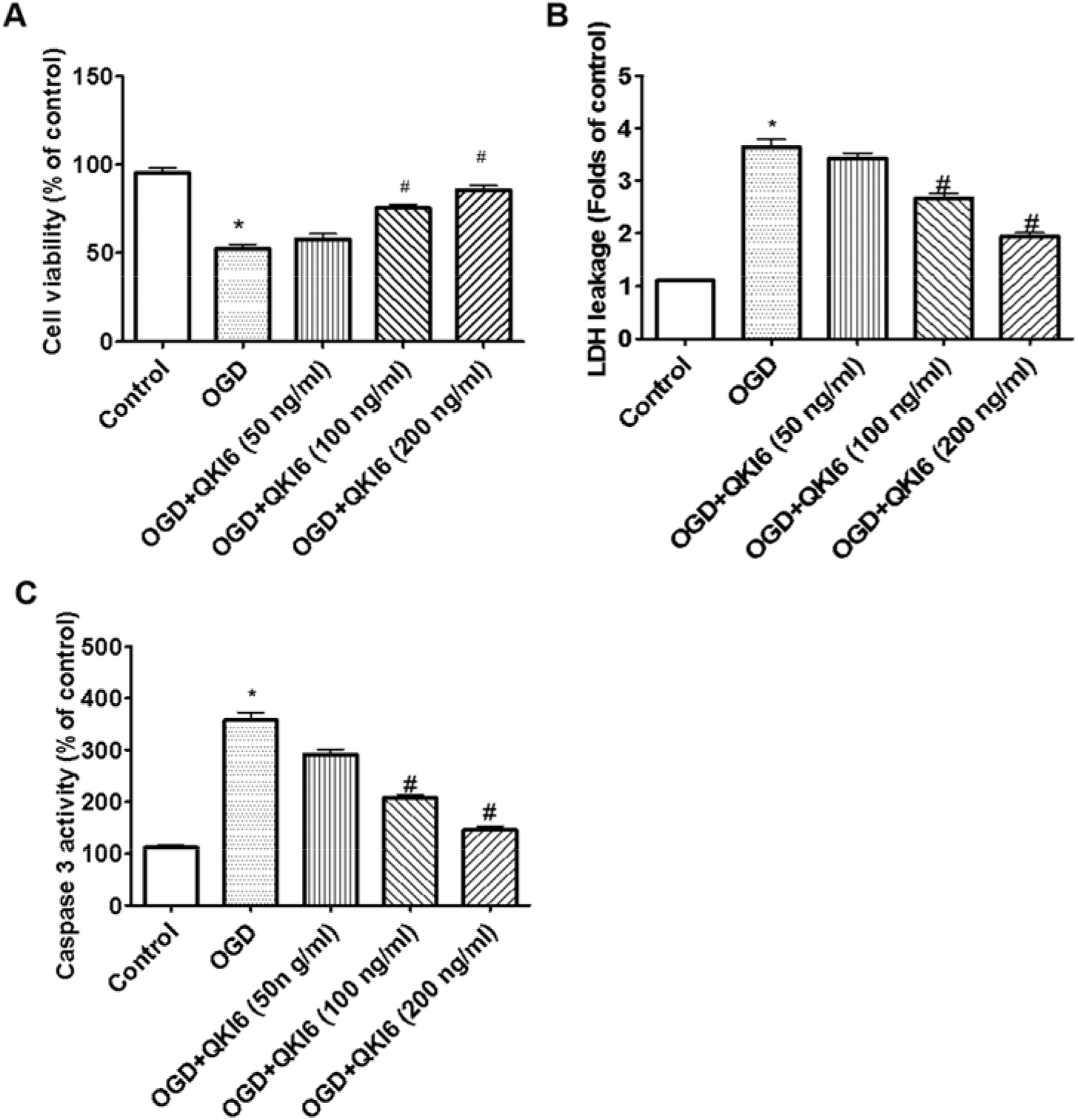
QKI 6 play a protective role in neuronal death come from OGD. The results of MTT experiment indicated that the viability of neurons after treatment with OGD obviously decreased, the survived cortical neurons was merely approximately 55%. However, when neurons were pretreated for 48 h with QKI 6 (50, 100, and 200 ng/mL, respectively) before 24h exposure to OGD. The viability assay demonstrated that neurons treated with QKI 6 was remarkably better than control neurons and dose dependence (*P*<0.01) (**A**). Further trial results form leakage rate of LDH assay confirmed that it increased obviously after OGD (**B**). These consequences figured out LDH release could be caused by OGD, which could be significantly inhibited by QKI 6 and dose dependence, too (*P*<0.001). Moreover, the protective roles of QKI 6 in apoptosis induced by OGD were evaluated with caspase-3 activity and TUNEL assays. The activity of caspase-3 obviously increased in cultured cortical neurons treated with OGD (**C**). But, the QKI 6 treatment could significantly suppressed the increased activity of caspase-3 in a dose-dependent manner (**C**). The data are presented as means ± SD from at six independent experiments. \**P*<0.01 (OGD group vs. Control group); #*P*<0.01 (vs. Control group).

Moreover, the protective roles of QKI 6 in apoptosis induced by OGD were evaluated with caspase-3 activity and TUNEL assays. The activity of caspase-3 obviously increased in cultured cortical neurons treated with OGD (**Fig. 1C**). Meanwhile, the staining of TUNEL increased remarkably in neurons treated with OGD (**Fig. 2**). But, the QKI 6 treatment could significantly suppressed the increased activity of caspase-3 (**Fig. 1C**) in a dose-dependent manner, and reduced the staining of TUNEL in neurons (**Fig. 2**). These results confirmed QKI 6 played the crucial protective role in apoptosis of neurons treated with OGD.

**Figure 2.**
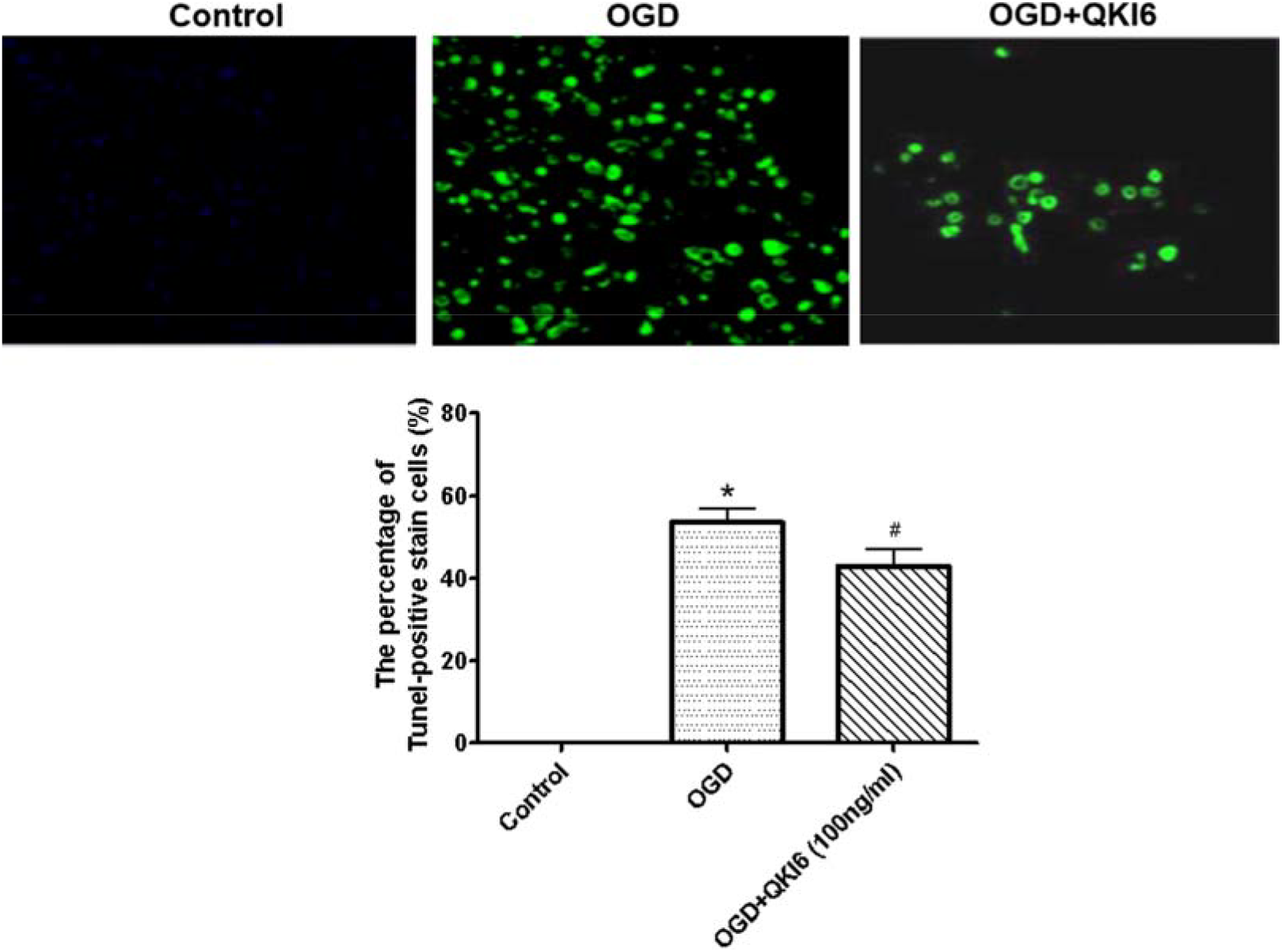
QKI 6 played a protective role in apoptosis of neurons treated with OGD with TUNEL assay. The staining of TUNEL increased remarkably in neurons treated with OGD. But, the QKI 6 treatment could significantly reduce the staining of TUNEL in neurons. These results confirmed QKI 6 played the protective role in apoptosis of neurons treated with OGD. The data are presented as means ± SD from at six independent experiments. \**P*<0.001 (OGD group vs. Control group); #*P*<0.01 (vs. Control group).

### The lipid metabolic disorders of primary cultured cortical neurons induced by OGD were reversed by QKI 6

Furthermore, the TGs (**Fig. 3A**) and cholesterol (**Fig. 3B**) content of primary cultured cortical neurons treated with OGD were significantly decreased compared with the controls, but could be restored by QKI 6. It demonstrated that the lipid metabolic disorders of primary cultured cortical neurons induced by OGD could be reversed by QKI 6 (**Fig. 3**).

**Figure 3.**
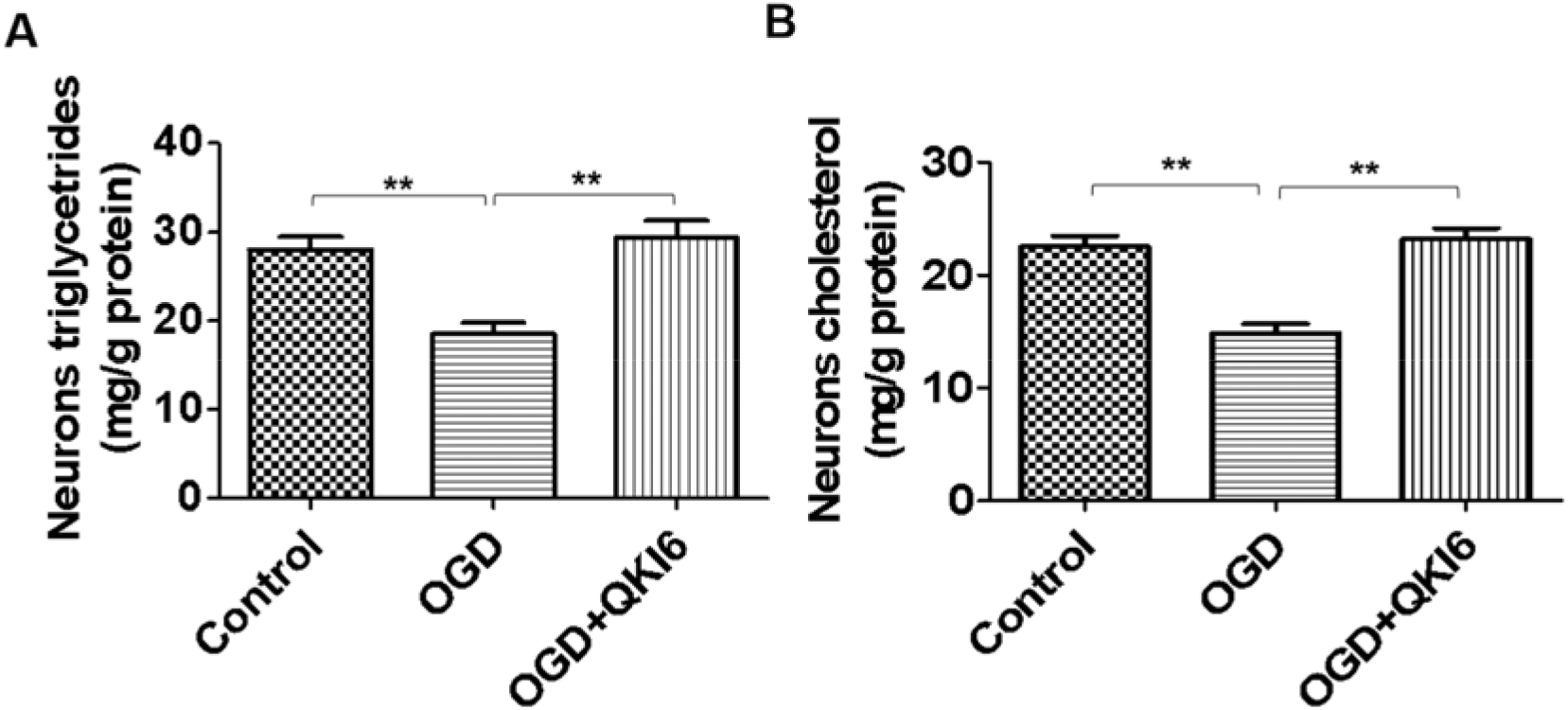
The lipid metabolic disorders of primary cultured cortical neurons induced by OGD are reversed by QKI 6. Furthermore, the TGs (**A**) and cholesterol (**B**) content of primary cultured cortical neurons treated with OGD were significantly decreased compared with the controls, but could be restored by QKI 6. It demonstrated that the lipid metabolic disorders of primary cultured cortical neurons induced by OGD could be reversed by QKI 6. The data are presented as means ± SD from at six independent experiments. \*\**P*<0.01.

### SIRT1 regulated the acetylation level of QKI 6

The acetylation level of QKI 6 increased in primary cultured cortical neurons induced by OGD, but it was reversed by SRT1720 (**Fig. 4A**). Additionally, the increasing acetylation level of QKI 6 was induced by inhibitor of SIRT1 (Niacinamide), but the QKI 6 acetylation decreased in primary cultured cortical neurons treated with OGD+SRT1720 (**Fig. 4B**). Furthermore, the increasing acetylation level of QKI 6 was induced by siRNA of SIRT1, but the QKI 6 acetylation decreased in primary cultured cortical neurons treated with Ad-SIRT1 (**Fig. 4C**). These results confirmed the protein interaction between SIRT1 and QKI 6, and its level in SRT1720 group was higher than in primary cultured cortical neurons induced by OGD (**Fig. 4D and 4E**).

**Figure 4.**
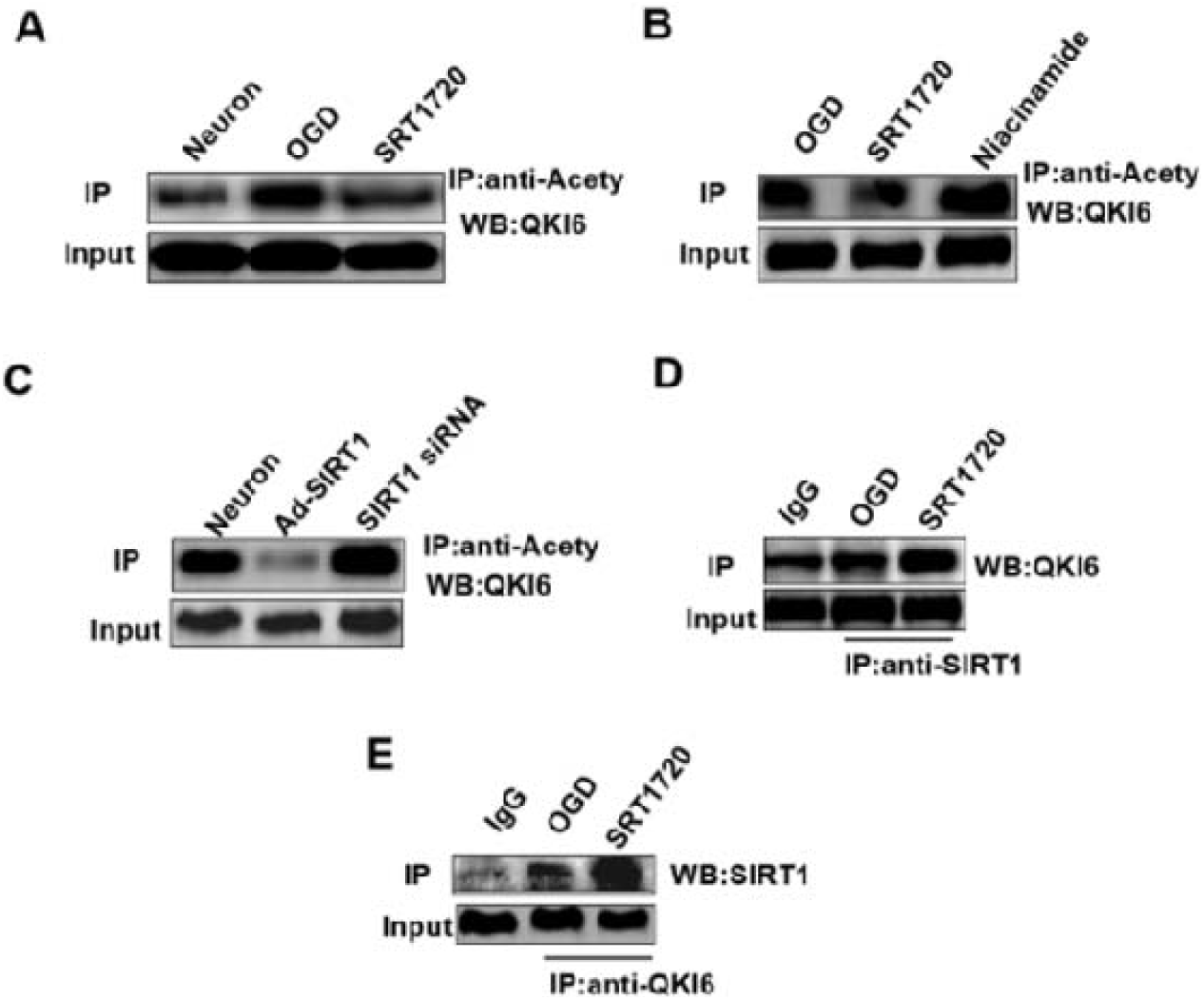
SIRT1 regulated the acetylation level of QKI 6. The acetylation level of QKI 6 increased in primary cultured cortical neurons induced by OGD, but it was reversed by SRT1720 (**A**). Additionally, the increasing acetylation level of QKI 6 was induced by inhibitor of SIRT1 (Niacinamide), but the QKI 6 acetylation decreased in primary cultured cortical neurons treated with OGD+SRT1720 (**B**). Furthermore, the increasing acetylation level of QKI 6 was induced by siRNA of SIRT1, but the QKI 6 acetylation decreased in primary cultured cortical neurons treated with Ad-SIRT1 (**C**). These results confirmed the protein interaction between SIRT1 and QKI 6, and its level in SRT1720 group was higher than in primary cultured cortical neurons induced by OGD (**D and E**). (The grouping of gels/blots had been sheared from different gels.)

### Neuronal SIRT1 regulated the synthesis of TGs in neurons via the QKI5/PPARγ/PGC-1α signaling pathway in vitro

To evaluate the ability of neuronal SIRT1 to maintain lipid homeostasis, the agonist (SRT1720) and inhibitor (Niacinamide) of SIRT1 was used in this study. Furthermore, adenovirus-mediated gene repletion of SIRT1 was employed in rat primary neurons, and siRNA of SIRT1 was used to interfere SIRT1 expression.

Furthermore, in primary neurons, SRT1720 upregulated SIRT1 expression, meanwhile expression of QKI 6, PGC-1α, and PPARγ increased, too. However, the SIRT1 expression level was reduced by Niacinamide, which also induced downregulation of QKI 6, PGC-1α, and PPARγ (**Fig. 5A**). But, SRT1720 induced downregulation of cleaved-caspase-3, cleaved-caspase-3 expression could be upregulated by Niacinamide (**Fig. 5A**). In addition, the decreased intracellular triglyceride content caused by SRT1720, whereas, Niacinamide enhanced the content of TG in primary neurons (**Fig. 5B**). Additionally, Ad-SIRT1 enhanced SIRT1, QKI 6, PGC-1α, and PPARγ expression in primary neurons, which was inhibited by SIRT1 siRNA (**Fig. 5C**). But, Ad-SIRT1 induced downregulation of cleaved-caspase-3, cleaved-caspase-3 expression could be upregulated by siRNA of SIRT1 (**Fig. 5C**). SRT1720 reduced the content of TGs in primary neurons, and which was promoted by SIRT1 siRNA (**Fig. 5D**). In addition, SIRT1 mediated the synthesis of TGs in neurons, which was associated with the QKI 6 and the PPARγ/PGC-1α signaling pathway.

**Figure 5.**
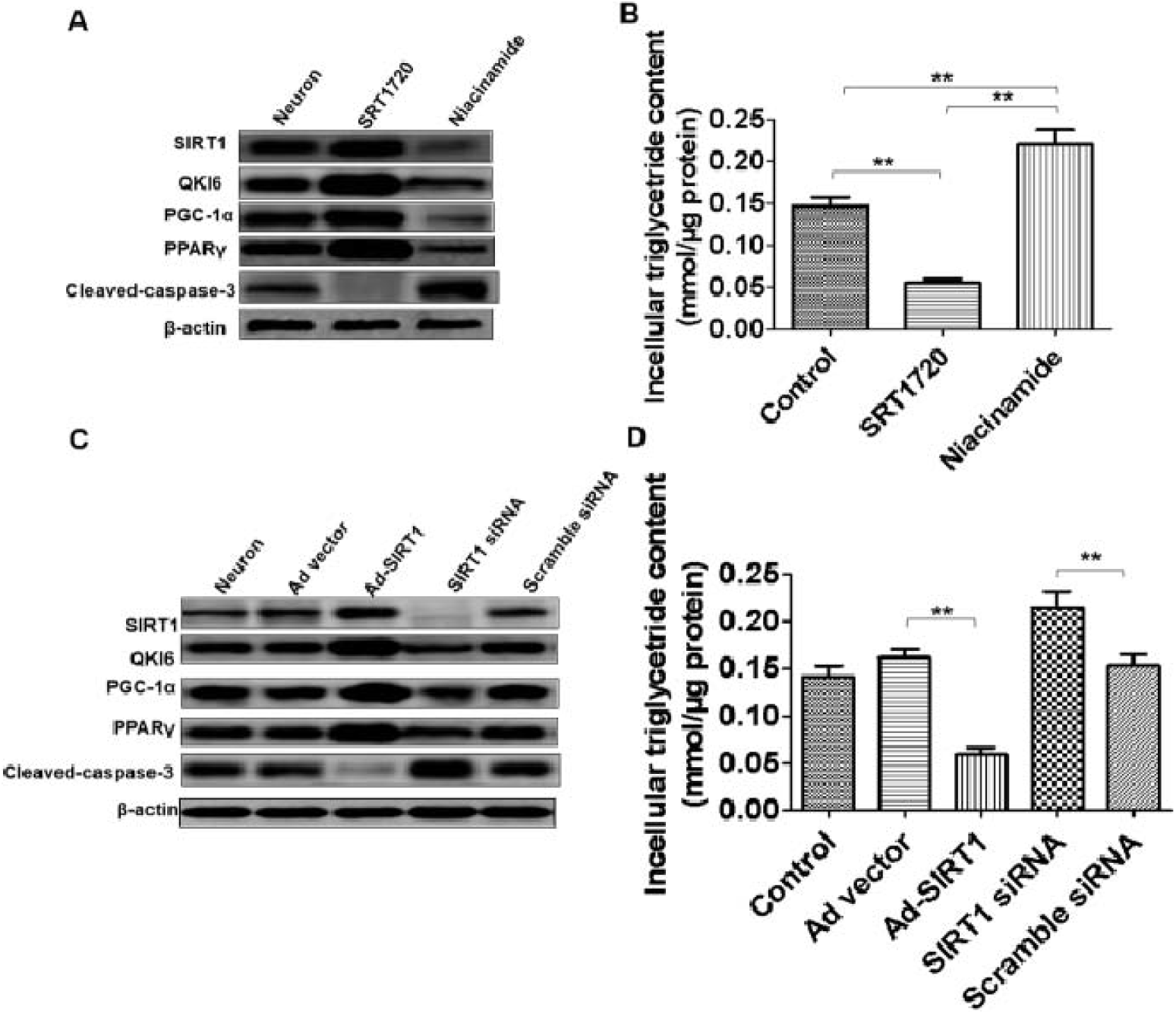
Neuronal SIRT1 regulated the synthesis of TGs in neurons via the QKI5/PPAR γ/PGC-1 α signaling pathway *in vitro*. In primary neurons, SRT1720 upregulated SIRT1 expression, meanwhile expression of QKI 6, PGC-1α, and PPARγ increased, too. However, the SIRT1 expression level was reduced by Niacinamide, which also induced downregulation of QKI 6, PGC-1α, and PPAR γ (**A**). But, SRT1720 induced downregulation of cleaved-caspase-3, cleaved-caspase-3 expression could be upregulated by Niacinamide (**A**). In addition, the decreased intracellular triglyceride content caused by SRT1720, whereas, Niacinamide enhanced the content of TG in primary neurons (**B**). Additionally, Ad-SIRT1 enhanced SIRT1, QKI 6, PGC-1α, and PPARγ expression in primary neurons, which was inhibited by SIRT1 siRNA (**C**). However, Ad-SIRT1 induced downregulation of cleaved-caspase-3, cleaved-caspase-3 expression could be upregulated by siRNA of SIRT1 (**C**). SRT1720 reduced the content of TGs in primary neurons, and which was promoted by SIRT1 siRNA (**D**). In addition, SIRT1 mediated the synthesis of TGs in neurons, which was associated with the QKI 6 and the PPARγ/PGC-1α signaling pathway. The data are presented as means ± SD from at six independent experiments. \*\**P*<0.01. (The grouping of gels/blots had been sheared from different gels.)

### Over-expression of QKI 6 protected against transient cerebral ischemia-induced secondary brain injury

Just after MCAO, the blood flow of these animal models (n=12) was determined, which decreased to 80-90% of the baseline level in the core area (**Fig. 6**), it was identified with successful. After MCAO, there was no significantly different of the cerebral blood flow between the two model groups. The representative brain sections showed total volume of infarction and infarction area in each group at 24 h after ischemia (**Fig. 7**). The total volume of infarction in Ad-QKI 6 group was 27.31±2.91 %, which was significantly smaller than those in vehicle group (60.36±6.75 %) (*P*<0.01) and Ad group (61.80±7.54 %) (*P*<0.01), respectively.

**Figure 6.**
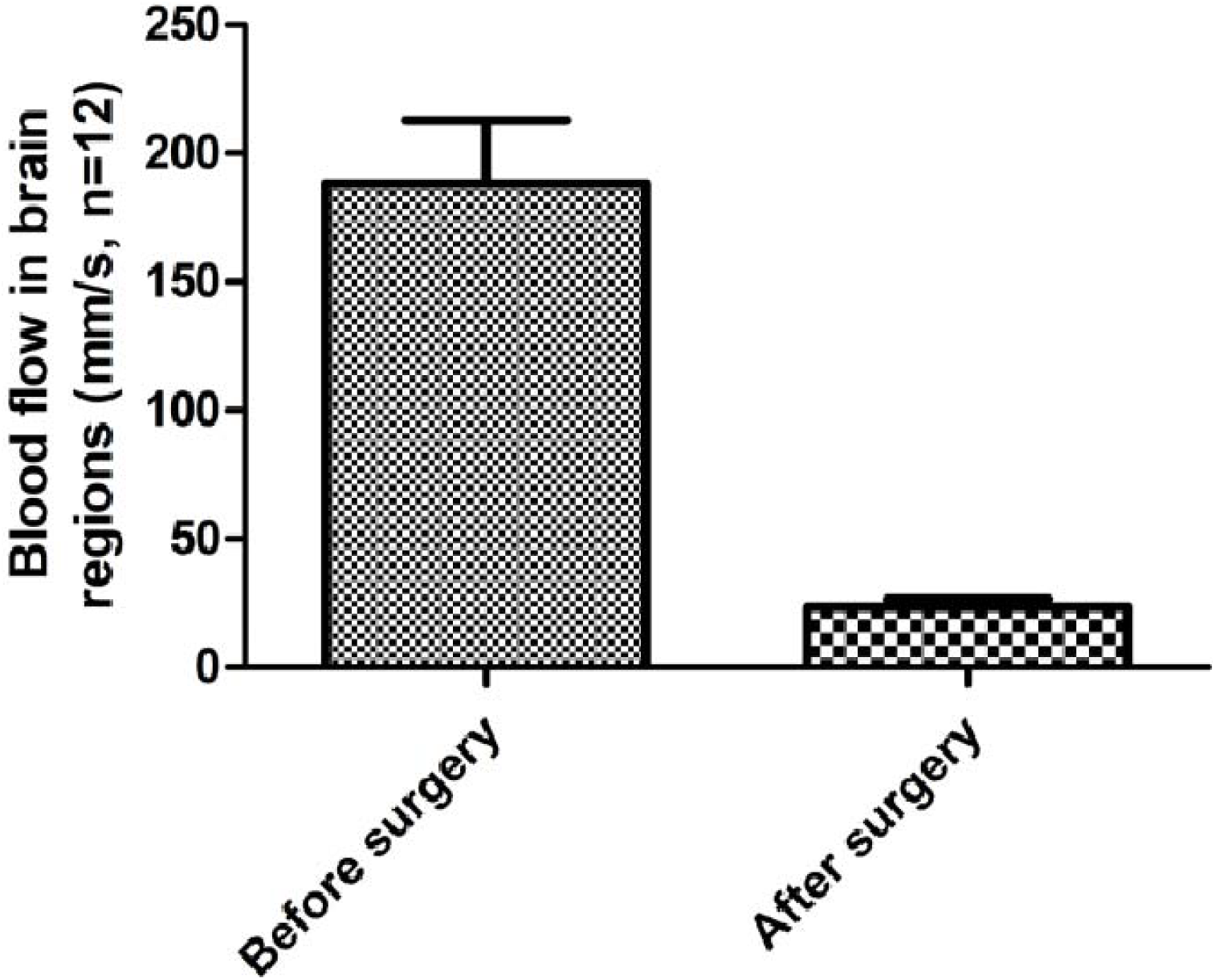
The level of blood flow in the core area. Just after MCAO, the blood flow of these animal models (n=12) was determined, which decreased to 80-90% of the baseline level in the core area.

**Figure 7.**
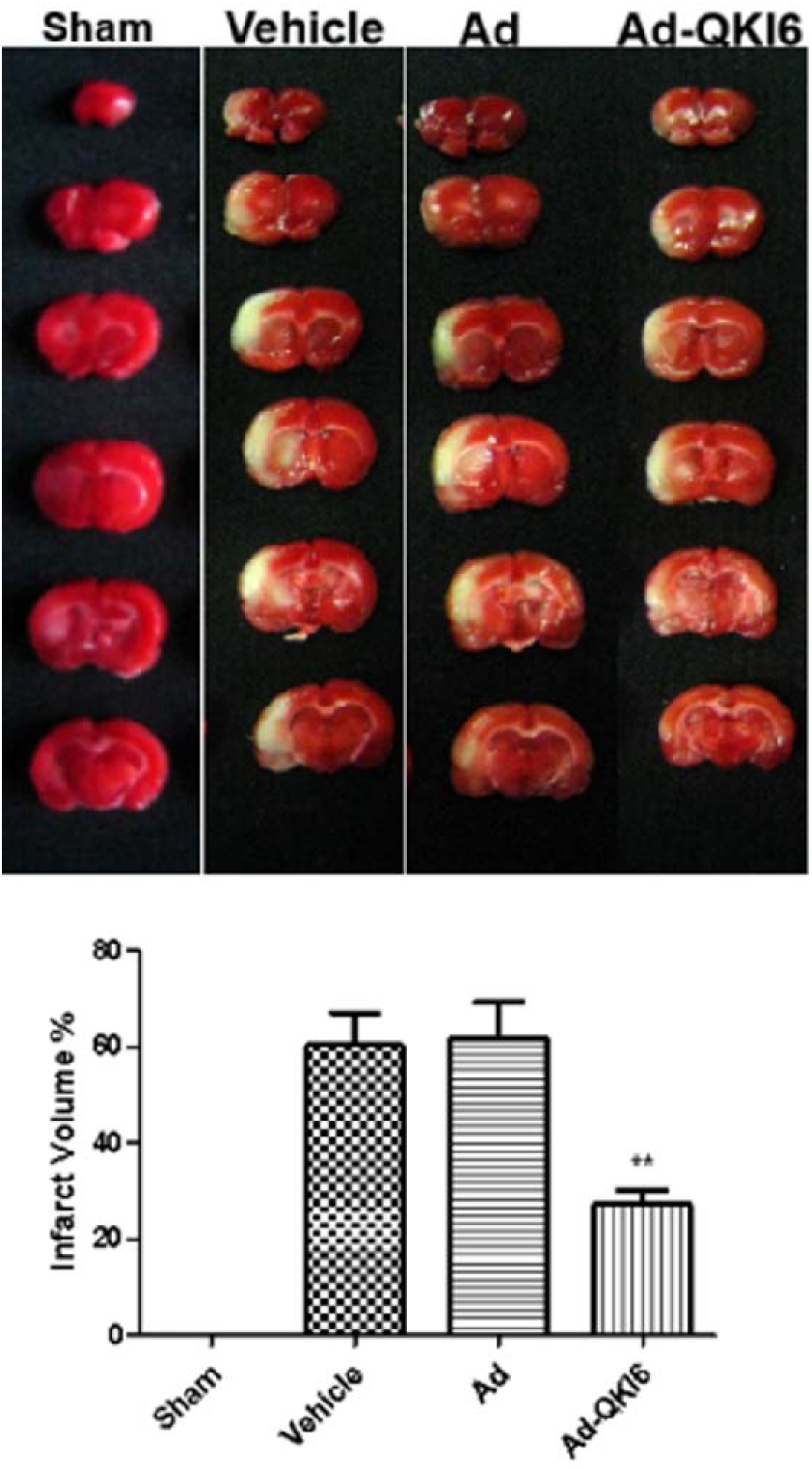
Over-expression of QKI 6 protected against transient cerebral ischemia-induced secondary brain injury. Just after MCAO, the blood flow of these animal models was determined, which decreased to 80-90% of the baseline level in the core area, it was identified with successful. After MCAO, there was no significantly different of the cerebral blood flow between the two model groups. The representative brain sections showed total volume of infarction and infarction area in each group at 24 h after ischemia. The total volume of infarction in Ad-QKI 6 group was 27.31±2.91 %, which was significantly smaller than those in vehicle group (60.36±6.75 %) (*P*<0.01) and Ad group (61.80±7.54 %) (*P*<0.01), respectively. The data are presented as means ± SD from at six independent experiments. \*\**P*<0.01 (Ad-QKI 6 group vs. Ad group or Vehicle group).

### Over-expression of QKI 6 with recombinant adenovirus reversed post-ischemic neuronal apoptosis induced by secondary brain damage in transient cerebral ischemia

In sham (**Fig. 8A**) and vehicle group (**Fig. 8B**), there was not any green fluorescent protein (GFP) expression. In contrast, expression of GFP was prevalent in the brain tissue (hippocampus, striatum, cortex penumbra) transfected with Ad (**Fig. 8C**) and Ad-QKI 6 (**Fig. 8D**), and the transfection rate was about 34.5±3.4 %. Then the expression level of QKI 6 was assessed in each group with western blot assay. In rats treated with QKI 6, the expression of QKI 6 increased significantly compared with that in Vehicle or Ad groups. In the tissues of cerebral cortex, which was obtained from the rats infected with recombinant adenovirus carrying QKI 6 gene, the expression of QKI 6 obviously increased (**Fig. 9A and 9B**). Transient cerebral ischemia injury induced an increased score of apoptotic index at 24 h after MCAO (**Fig. 9C**). Over expression of QKI 6 reversed the apoptosis score compared to the group with transient cerebral ischemia injury. The apoptotic index of Ad-QKI 6 group (18.28±0.18) % was remarkably lower than Ad group (57.68±2.11) and Vehicle group (58.22±1.96) % (*P*<0.001), respectively. The final ordinary pathway of caspase-dependent apoptosis, caspase-3 activity of neuron, neuronal apoptosis induced by ischemic was further examined after transfection with QKI 6 gene to identify its specific role. The caspase-3 activation in brains induced by ischemia/reperfusion reduced remarkably after treatment with QKI 6 gene (**Fig. 9D**). In brains of sham group, the percentage of detected positive cells staining with TUNEL was 0.1%. It demonstrated that the detectable neuronal apoptosis could not be caused by surgical procedure. Conversely, in the tissues from brains of ischemic-reperfused rats, the nuclei with TUNEL-positive stain were prevalent. But, it were reduced significantly through treatment with QKI 6 gene (**Fig. 10**). QKI 6 gene over-expression might play an prohibitive role in post-ischemic neuronal apoptosis.

**Figure 8.**
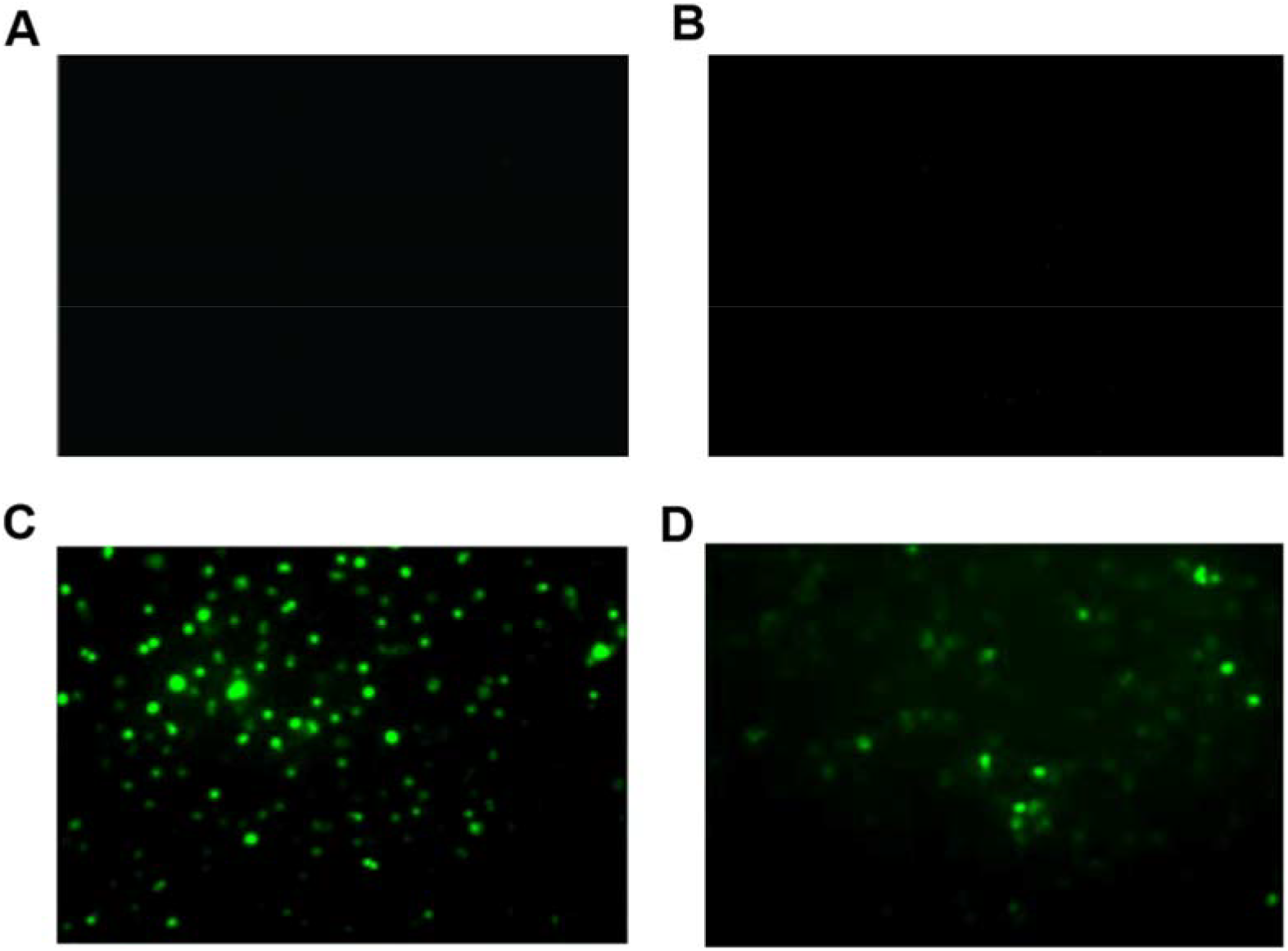
Over-expression of QKI 6 induced with recombinant adenovirus. In sham (**A**) and vehicle group (**B**), there was not any green fluorescent protein (GFP) expression. In contrast, expression of GFP was prevalent in the brain tissue (hippocampus, striatum, cortex penumbra) transfected with Ad (**C**) and Ad-QKI 6 (**D**), and the transfection rate was about 34.5±3.4 %.

**Figure 9.**
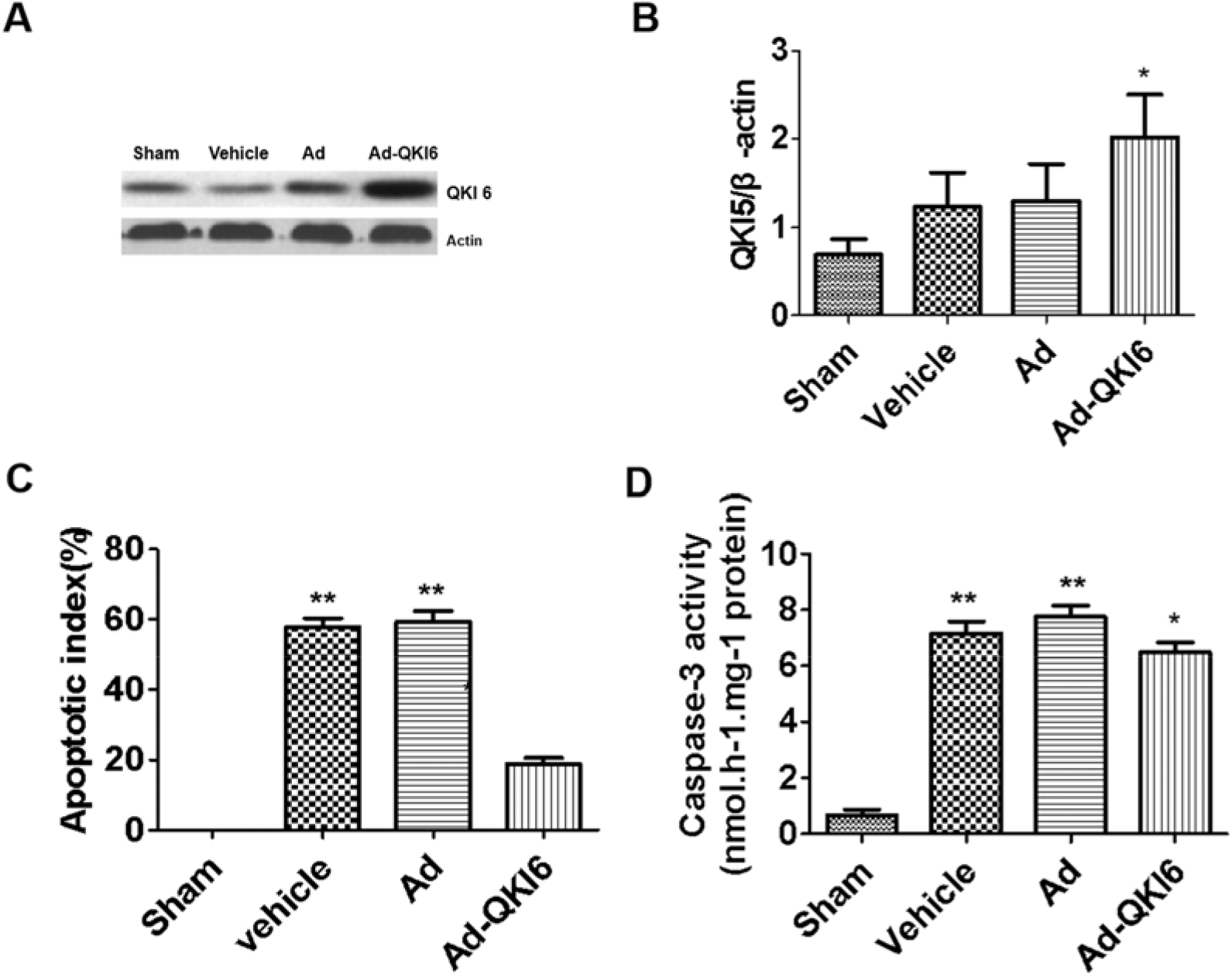
Over-expression of QKI 6 with recombinant adenovirus reversed post-ischemic neuronal apoptosis caused by secondary brain damage in transient cerebral ischemia. Then the expression level of QKI 6 was assessed in each group with western blot assay. In rats treated with QKI 6, the expression of QKI 6 increased significantly compared with that in Vehicle and Ad groups. In the tissues of cerebral cortex, which was obtained from the rats infected with recombinant adenovirus carrying QKI 6 gene, the expression of QKI 6 (**A and B**) obviously increased. Transient cerebral ischemia injury induced an increased score of apoptotic index (**C**) at 24 h after MCAO. Over expression of QKI 6 reversed the apoptosis score compared to the group with transient cerebral ischemia injury. The apoptotic index of Ad-QKI 6 group (18.28±0.18) % was remarkably lower than Ad group (57.68±2.11) and Vehicle group (58.22±1.96) % (*P*<0.001), respectively. The final ordinary pathway of caspase-dependent apoptosis, caspase-3 activity of neuron, neuronal apoptosis induced by ischemic was further examined after transfection with QKI 6 gene to identify its specific role. The caspase-3 activation in brains induced by ischemia/reperfusion reduced remarkably after treatment with QKI 6 gene (**D**). The data are presented as means ± SD from at six independent experiments. \**P*<0.01 (Ad group or Vehicle group vs. Sham group) ;\**P*<0.01 (Ad-QKI 6 group vs. Ad group or Vehicle group) . (The grouping of gels/blots had been sheared from different gels.)

**Figure 10.**
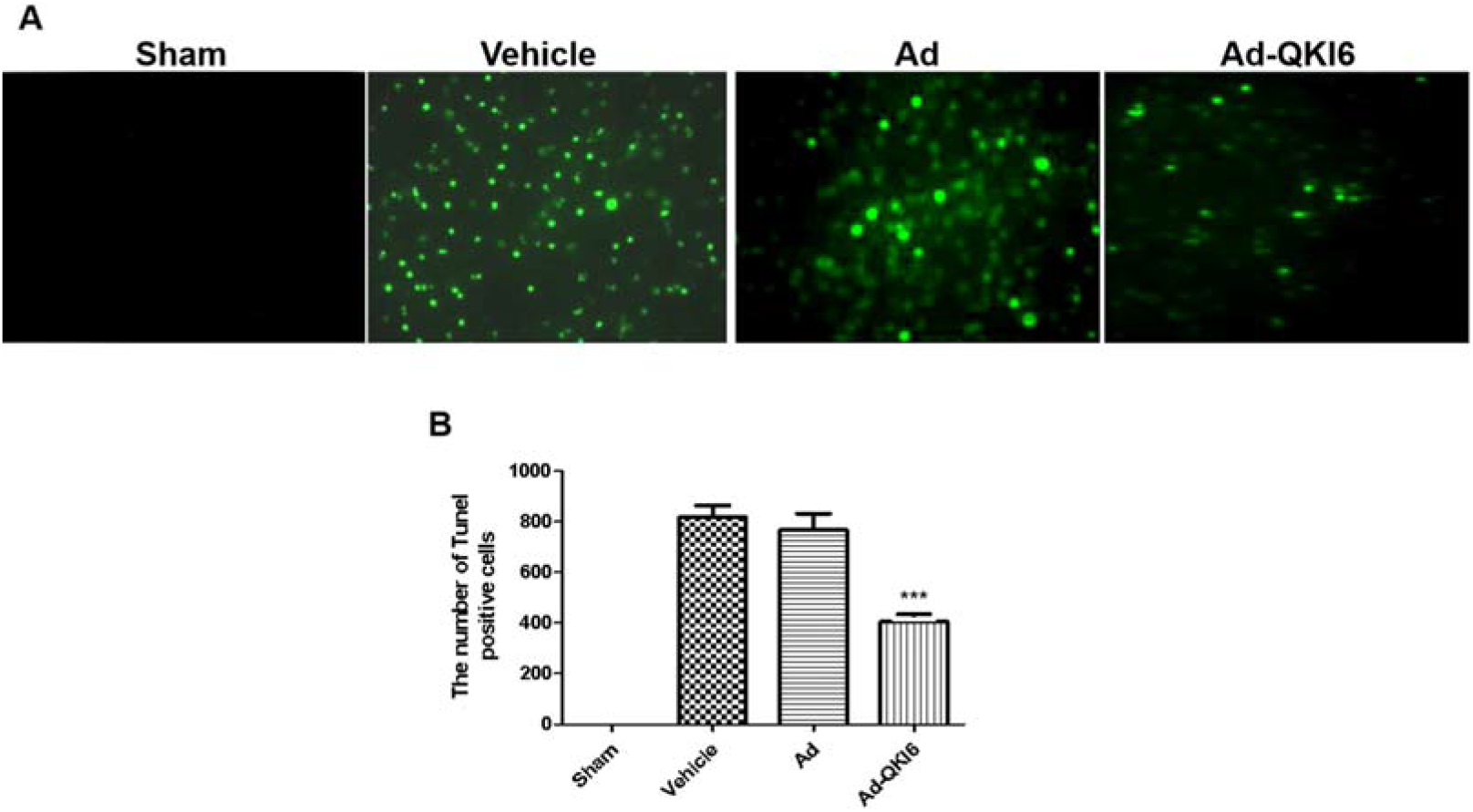
Over-expression of QKI 6 with recombinant adenovirus reversed post-ischemic neuronal apoptosis (TUNEL assay) induced by secondary brain damage in transient cerebral ischemia. In brains of sham group, the percentage of detected positive cells staining with TUNEL was 0.1%. It demonstrated that the detectable neuronal apoptosis could not be caused by surgical procedure. Conversely, in the tissues from brains of ischemic-reperfused rats, the nuclei with TUNEL-positive stain were prevalent. But, it were reduced significantly through treatment with QKI 6 gene. QKI 6 gene over-expression might play an prohibitive role in post-ischemic neuronal apoptosis.

### Over-expression of QKI 6 with recombinant adenovirus reverses behavioral impairments induced by secondary brain damage in transient cerebral ischemia via PPARγ/PGC-1α signaling pathway

Transient cerebral ischemia damage could reduce the neurological score (**Fig. 11**) at 24 h after MCAO. Over-expression of QKI 6 may reverse the score compared to transient cerebral ischemia injury group. The neurological score of Ad-QKI 6 group (12.45±0.18 %) were higher than that in Ad group (9.96±0.09%) and Vehicle group (9.82±0.17%) (*P*<0.001), respectively. Over-expression of QKI 6 gene played a potential protective role against secondary brain damage of cerebral cortex induced by transient cerebral ischemia after MCAO, additionally, may restore neurological/behavioral impairment.

**Figure 11.**
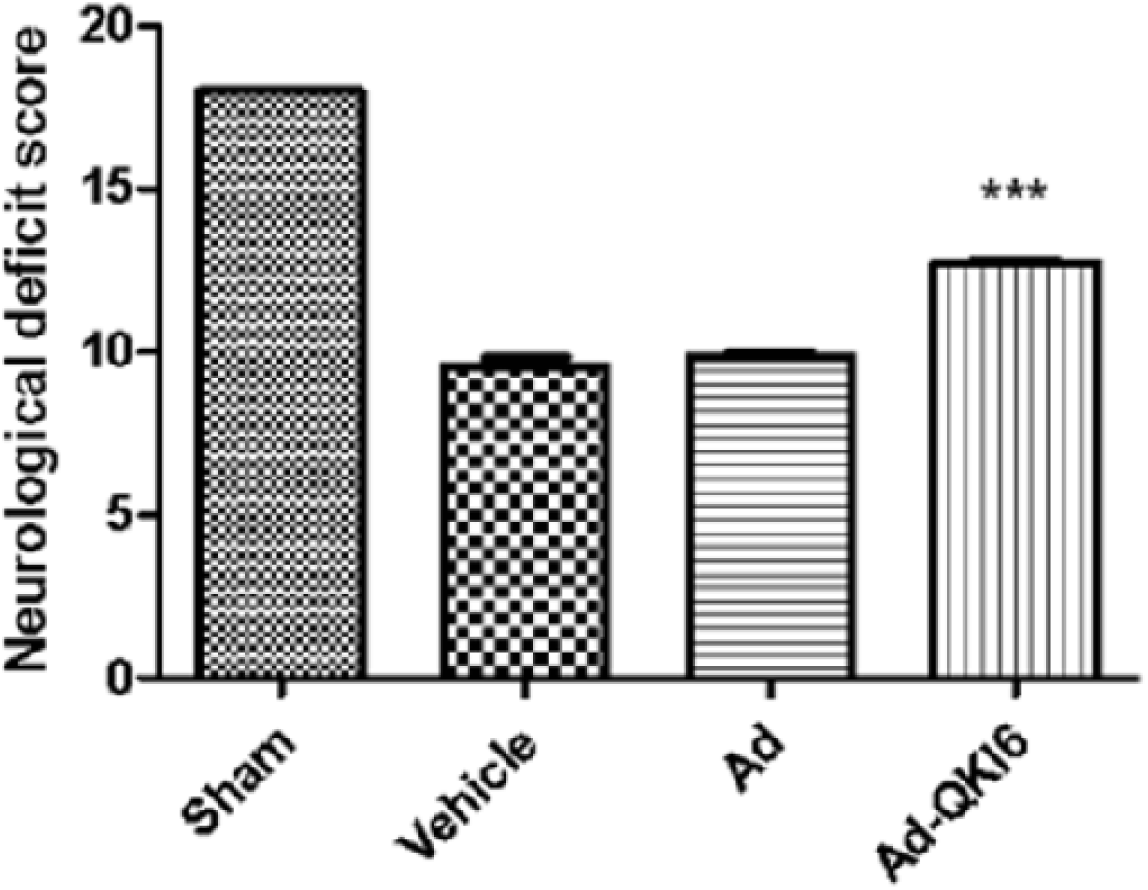
Over-expression of QKI 6 with recombinant adenovirus reversed behavioral impairments induced by secondary brain damage in transient cerebral ischemia. Transient cerebral ischemia damage could reduce the neurological score at 24 h after MCAO. Over-expression of QKI 6 may reverse the score compared to transient cerebral ischemia injury group. The neurological score of Ad-QKI 6 group (12.45±0.18 %) were higher than that in Ad group (9.96±0.09%) and Vehicle group (9.82±0.17%) (*P*<0.001), respectively. Over-expression of QKI 6 gene played a potential protective role against secondary brain damage of cerebral cortex induced by transient cerebral ischemia after MCAO, meanwhile may restore neurological/behavioral impairment. The data are presented as means ± SD from at six independent experiments. \*\*\**P*<0.000 (Ad-QKI 6 group vs. Ad group or Vehicle group) .

The results of western blot assay demonstrated that the expression level of SIRT1, QKI 6, PPARγ, and PGC-1 α decreased in model rat, but it was reversed by QKI 6 (**Fig. 12**). However, over-expression of QKI 6 with its recombinant adenovirus induced downregulation of cleaved-caspase-3 in CIRI model come from secondary brain damage of transient cerebral ischemia. It demonstrated that neuronal SIRT1 regulated expression of QKI 6 via SIRT1-PPARγ-PGC-1 α signaling pathway axis. Besides, which was related to activated caspase-3 (cleaved-caspase-3) and apoptosis.

**Figure 12.**
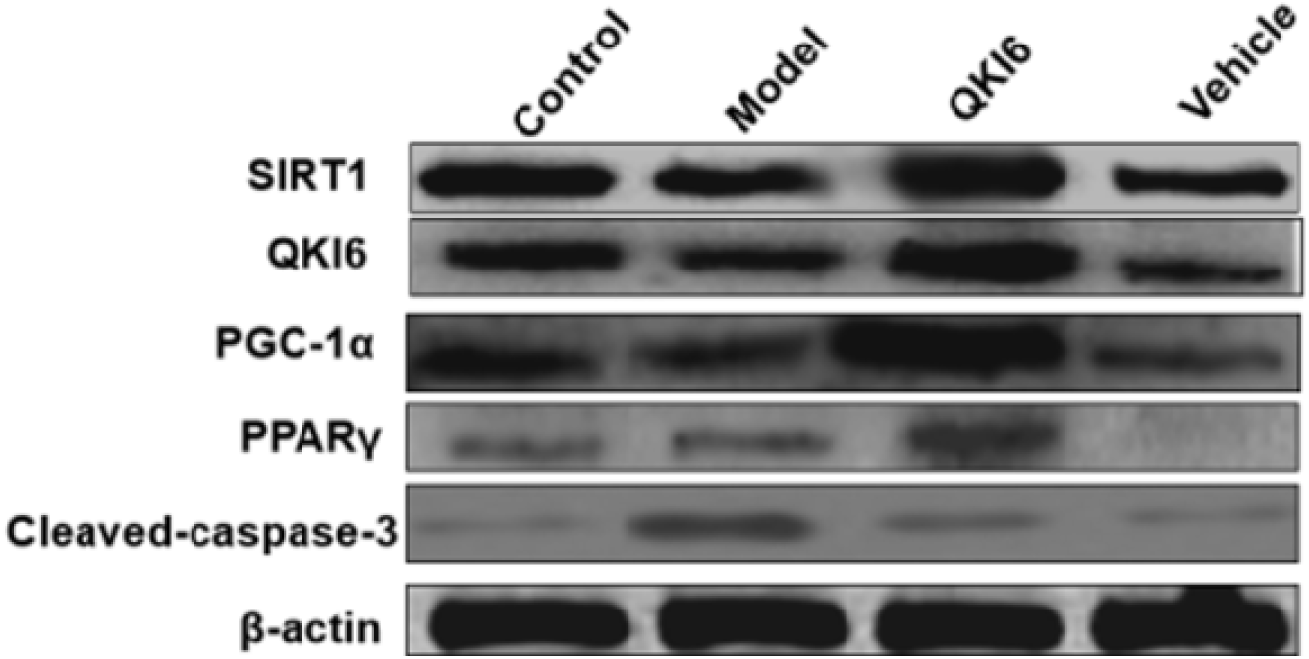
Over-expression of QKI 6 with recombinant adenovirus reversed CIRI induced by secondary brain damage in transient cerebral ischemia via PPAR γ /PGC-1α signaling pathway. The results of western blot assay demonstrated that the expression level of SIRT1, QKI 6, PPARγ, and PGC-1 α decreased in model rat, but it was reversed by QKI 6. It demonstrated that neuronal SIRT1 regulated expression of QKI 6 via SIRT1-PPARγ -PGC-1 α signaling pathway axis. However, over-expression of OKI5 with its recombinant adenovirus induced downregulation of cleaved-caspase-3 in CIRI model come from secondary brain damage of transient cerebral ischemia. (The grouping of gels/blots had been sheared from different gels.)

## Disscusion

RNA-binding protein-mediated post-transcriptional regulation is rarely studied in neuronal metabolism. STAR family member QKI is an RNA-binding protein that produces multiple isoforms in human body, and showed a variety of expression pattern in various kinds of tissue cells [28]. Among them, QKI 6 is mainly located in the nucleus, but can also shuttle to the cytoplasm [29, 30], while QKI 6 and QKI 7 are mainly distributed in the cytoplasm. By binding to the specific recognition element (QRE) of the 3’UTR of mRNA [31, 32], QKI plays a key role in regulating the cytoplasmic/nuclear localization, stability, and translation efficiency of mRNA [33, 34]. We had found that QKI is expressed in primary neurons extracted from cerebral cortex of rat, and the mainly expressed isomer was QKI 6, but its functional studies in the brain had not been reported.

SIRT1 is mediated by post-translational regulation, and many transcription factors such as FOXO1, SREBP, ChREBP, PPARα, PPARγ, and PGC1α are mediated through SIRT1 at the transcriptional level [35]. QKI is an RNA-binding protein whose mediated regulation is post-transcriptional regulation of RNA levels. Specific to RNA sequence binding, QKI is widely involved in variable splicing, subcellular localization, stability maintenance, and translational regulation of RNA.

The major reasons of death and long-term disability worldwide include stroke. It is related to obvious socioeconomic and clinical implications, especially need effective therapies. In practice, stroke is a pathological process in which multi-factors are involved, mainly related to oxidative stress, inflammation, calcium overload, brain edema and cell apoptosis and other relevant events [36]. TIAs (transient ischemic attacks) are known as brief ischemic episodes according to clinical practice, and for over two decades have been studied [37, 38]. The tolerance could be induce by TIAs through elevating the threshold of human brain tissue vulnerability [39]. It is the critical response for neuroprotection and the potential molecular mechanisms. Till now, there are no clinically effective therapies for stroke, in spite of a great number of studies based on experimental neuroprotective compounds and animal model demonstrated the promising results. To investigate potential molecular mechanisms associated with TIAs may discover the novel targets on the prevention and treatment for stroke [40, 41].

Therefore, rat neurons were firstly isolated in our study and used to characterize the roles of function and molecular mechanisms of QKI 6, whether QKI 6 protects against neuronal death induced by OGD. OGD induced remarkably LDH release of neurons, but neurons treated with QKI 6 followed with OGD were remarkably better than control neurons only treated with OGD. The various dose of QKI 6 suppressed obviously LDH release with dose dependence. Moreover, QKI 6 provided significantly protective effects on neuronal apoptosis induced by OGD. Which was identified by the results of caspase-3 activity and TUNEL assays. Meanwhile, the lipid metabolic disorders of primary cultured cortical neurons induced by OGD could be reversed by QKI 6.

Further investigation had been conducted to confirm the molecular mechanism of QKI 6. The online local tool of KA-predictor (http://sourceforge.net/p/ka-predictor) was used to predict the acetylation modification site of QKI 6. Therefore, SIRT1 was predicted that it may acetylate QKI 6. We found that the acetylation level of QKI 6 increased in primary cultured cortical neurons induced by OGD, but it was reversed by SRT1720 (the activator of SIRT1). Additionally, the acetylation level of QKI 6 increased in primary cultured cortical neurons induced by OGD, but it was reversed by SRT1720. Additionally, the increasing acetylation level of QKI 6 was induced by inhibitor of SIRT1 (Niacinamide), but the QKI 6 acetylation decreased in primary cultured cortical neurons treated with OGD+SRT1720. Furthermore, the increasing acetylation level of QKI 6 was induced by siRNA of SIRT1, but the QKI 6 acetylation decreased in primary cultured cortical neurons treated with Ad-SIRT1. These results confirmed the protein interaction between SIRT1 and QKI 6, and its level in SRT1720 group was higher than in primary cultured cortical neurons induced by OGD.

To evaluate the ability of neuronal SIRT1 to maintain lipid homeostasis, the agonist (SRT1720) and inhibitor (Niacinamide) of SIRT1 was used in this study. Furthermore, adenovirus-mediated gene repletion of SIRT1 was employed in rat primary neurons, and siRNA of SIRT1 was used to suppress SIRT1 expression. These experiments identified that in primary neurons, SIRT1 mediates the synthesis of TGs in OGD model, which is associated with the QKI 6 and the PPARγ/PGC-1α signaling pathway. Besides, the MCAO-induced CIRI rat model was well established and used to characterize the effects of QKI 6 mediated by SIRT1 on synthesis of triglyceride in neuron and neuronal apoptosis via activation of SIRT1-PPARγ-PGC-1α signaling pathway. It demonstrated that over-expression of QKI 6 with recombinant adenovirus protected against transient cerebral ischemia-induced by secondary brain injury, reversed post-ischemic neuronal apoptosis and behavioral impairments induced by secondary brain damage in transient cerebral ischemia via PPARγ/PGC-1 α signaling pathway.

## Conclusions

Our results demonstrated SIRT1 deacetylates QKI 6, the RNA-binding protein, that affects significantly the synthesis of triglyceride in neurons of CIRI rat model. Moreover, it activated transcription factor PGC-1 α through post-transcriptional regulation of the expression of PPARγ, and further enhanced synthesis of triglyceride, thereby restrained the progression of neural apoptosis and CIRI.

## Acknowledgment

None.

## Financial competing interests

### Funding

This study was funded by National Natural Science Foundation of China (No. 81771469), Shanxi Province Social Development Science and Technology Attack Project (No. 2016SF-132), Logistics Research Program of Chinese People’s Liberation Army (No. CJN14J005) and Science and Technology Project of Louyang, Henan Province, China (No. 1603002A-13).

### Employment

None.

### Personal financial interests

None.

### Ethical approval

All applicable international, national, and institutional guidelines for the care and use of animals were followed.

### Informed consent

None.

### Author Contribution

RL and HZL contributed to the conception of the study. RL, JYD and QQW contributed significantly to analysis and manuscript preparation; CHL, QX, and QC performed the data analyses and wrote the manuscript; RL and QC helped perform the analysis with constructive discussions.

